# Establishing conditions for the generation and maintenance of estrogen receptor-positive organoid models of breast cancer

**DOI:** 10.1101/2023.08.09.552657

**Authors:** Michael UJ Oliphant, Dipikaa Akshinthala, Senthil K Muthuswamy

**Affiliations:** Department of Cell Biology and Ludwig Cancer Center, Harvard Medical School, Boston, MA, USA; Laboratory of Cancer Biology and Genetics, Center For Cancer Research, National Cancer Institute, National Institute of Health, Bethesda, MD, USA

## Abstract

Patient-derived organoid models of estrogen receptor-positive (ER+) breast cancer would provide a much-needed tool to better understand drug resistance and disease progression. However, the establishment and long-term maintenance of ER expression, function, and response in vitro remains a significant challenge. Here, we report the development of an ER+ breast tumor organoid medium (BTOM-ER) that conserves ER expression, estrogen responsiveness, and dependence, as well as sensitivity to endocrine therapy of ER+ patient-derived xenograft organoids (PDXO). Our findings demonstrate the utility of subtype-specific culture conditions that better mimic the characteristics of the breast epithelial biology and microenvironment, providing a powerful platform for investigating therapy response and disease progression of ER+ breast cancer.

## Introduction

Breast cancer remains one of the most frequently diagnosed cancers and cause of cancer deaths among women worldwide (Siegel et al., 2022). Despite estrogen receptor-positive (ER+) breast cancer being the most prevalent subtype, the establishment of clinically relevant models remains a challenge. More recently, three-dimensional (3D) organoid models established from patient tumors bridge the gap between cell culture and patient-derived xenograft platforms (Dekkers et al., 2021; Guillen et al., 2022; Sachs et al., 2018). Previous studies have shown that organoid media formulations are critical for the establishment, growth, and maintenance of patient-specific characteristics of ER+ breast cancer organoid models(Guillen et al., 2022; Hogstrom et al., 2023; Sachs et al., 2018). However, consistent establishment, decreased estrogen receptor (ER) expression and loss of ER-dependent transcription after extended culturing continues to be a challenge for the generation and use of organoid models from ER+ breast tumors.

Here we report the development of a simplified media considering breast tissue-relevant cytokines and growth factors that support the establishment and expansion of organoids from ER+ breast tumors. The conditions reported here support both retention of ER expression over long-term culture, responsiveness to estrogen, and sensitivity to the anti-hormone therapeutic agent, fulvestrant, identifying a new approach for generating ER+ organoid models for breast cancer.

## Results

### Optimization of media conditions that support robust growth of ER+ PDX Organoids

In designing the media, serum was excluded because of its undefined nature and to prevent the proliferation of fibroblasts. We also omitted using stem cell factors to prevent reprogramming or the expansion of undifferentiated subpopulations of cells at the expense of differentiated epithelial cells. For instance, R-spondin-1 was not included due to its roles in promoting stem and basal epithelial cell differentiation and proliferation (Broutier et al., 2016; Huch et al., 2015; Jardé et al., 2016). Noggin was not included for its role in promoting stemness due to evidence that it can interfere with ER transcriptional responses (Páez-Pereda et al., 2003). We included bovine pituitary extract (BPE) and B27 as a serum-free mitogenic supplement (Hammond et al., 1984; Maciag et al., 1981) and to protect against oxidative stress, a common challenge that occurs in 3D cultures (Kent & Bomser, 2003). To identify the growth factors needed, we reviewed growth factors and nutrients that are known to play critical roles in the growth and maintenance of normal ER+ breast epithelial cells and those implicated in ER biology in developmental biology studies. Prolactin stimulates ER expression and works in concert with progesterone receptor to promote autocrine secretion-mediated mammary epithelial cell and breast cancer proliferation (Frasor & Gibori, 2003; Gutzman et al., 2004;Naylor et al., 2003). Amphiregulin supports ER expression and estrogen signaling in the mammary gland (Ciarloni et al., 2007a, 2007b). IL6 expression is positively correlated with hormone receptor-positive tumors and has been shown to coordinate estrogen expression in vivo (Fontanini et al., 1999; Martínez-Pérez et al., 2021; Purohit et al., 1995, 2002; Sasser et al., 2007; Speirs et al., 2000). The above factors were added to DMEM/F12 media in conjunction with growth factors frequently used to support the proliferation of mammary epithelial cells in culture and in vivo, including FGF2, FGF10, EGF, insulin, and hydrocortisone (See Suppl.1 for details of media recipe and preparation). Supplementation of estradiol to the media caused varying effects on cell growth; some organoid cultures showed no difference, whereas others showed a non-significant increase in growth (Suppl. Fig. 1). Furthermore, estrogen receptor expression and activity can be negatively regulated in both normal and cancer contexts during prolonged exposure to E2 in culture and in vivo (Bondar et al., 2009; Borrás et al., 1994; Hatsumi & Yamamuro, 2006); hence no E2 supplementation was used for routine culture.

We used two independent ER+ patient-derived xenograft tumors (Guillen et al., 2022) to find the optimal concentrations of the various growth factors to define a Breast Tumor Organoid Media for ER+ breast cancers (BTOM-ER) that supports growth and expansion (Fig 1a, Table 1). The PDXOs achieved robust growth with an average doubling time of 2.042 days (Fig 1b) and remained stable in later passages. The phase and histomorphology of PDXOs were heterogeneous within cultures consisting of solid and hollow organoids with the formation of budding structures between 10-14 days post-plating (Fig 1c). Both proliferation rates and phase morphological characteristics were relatively consistent over 15 passages, as determined by phase morphology and Incucyte live-cell imaging (data not shown). Importantly, immunofluorescence analysis for cytokeratin 8/18 and estrogen receptor demonstrated the presence of ER+ luminal tumor epithelia (Fig 1d). Collectively, these data suggest that BTOM-ER allows for the establishment and maintenance of ER+ luminal epithelial organoid cultures that maintain proliferative capacity during long-term culture.

**Figure 1.**
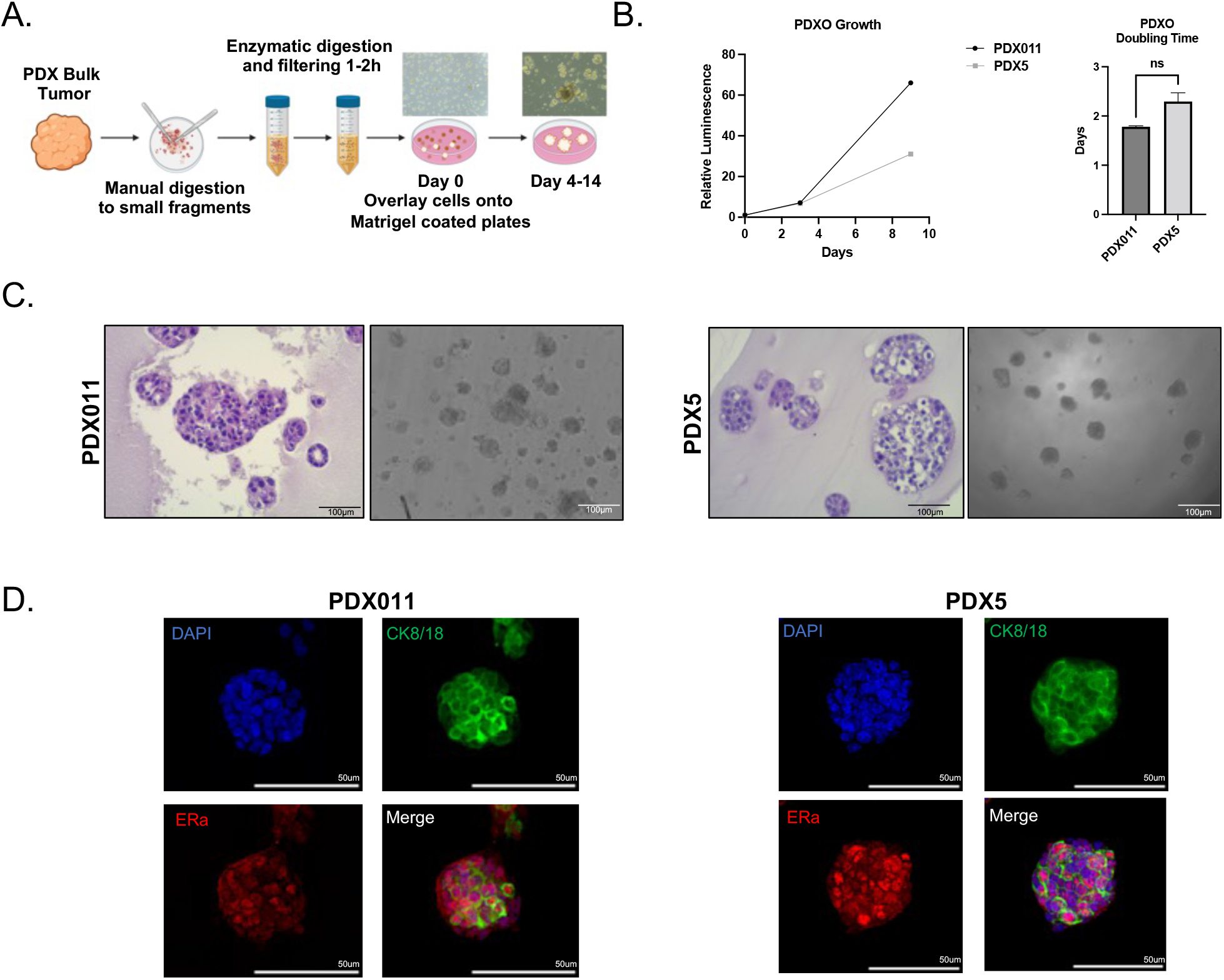
BTOM promotes robust growth and ER expression of ER+ PDXOs. A) Schematic describing establishment of PDXOs. B) A representative experiment of growth over time of PDX011 and PDX5 organoids, as measured by 3D Cell Titer Glo (Promega). Doubling time was calculated using GraphPad Prism Software. C) H&E and brightfield images to show the morphology of established PDXOs. Scale bars: 100um D). Immunofluorescence of PDXOs displaying CK8/18 and ER-alpha expression. DAPI was used to stain nuclei.

**Table 1.** PDXO Clinical Information.

| **Patient ID** | **ER** | **PR** | **HER2** | **ER Mutation Status** | **Additional genetic alterations** | **Morphology** |
| --- | --- | --- | --- | --- | --- | --- |
| PDX011 | + | + | - | WT | N/A | Solid spheres, with branching for larger organoids |
| PDX5 | + | + | + | L536P | RBM10 and RNF111 deletions | Solid spheres with small numbers of protruding cells |

**Table 2:** Components of BTOM-ER growth media.

|  | **Supplements/Growth factors** | **Vendor** | **Catalog #** | **Final concentration** |
| --- | --- | --- | --- | --- |
| **Reagent A** |  |  |  |  |
| 1 | Bovine Pituitary Extract | Hammond cell tech | 1078-NZ | 0.8 ml for 100 ml |
| 2 | B27 | Thermo | 17504001 | 1.0 ml for 100 ml |
| 3 | Recombinant Human FGF- Basic (FGF2) | Peprotech | AF-100- 18B | 10 ng/ml |
| 4 | Recombinant Human FGF10 | Peprotech | 100-26 | 10 ng/ml |
| 5 | Recombinant Human EGF | Peprotech | AF-100-15 | 2.0 ng/ml |
| 6 | Recombinant Human IL6 | Peprotech | 200-06 | 100 ng/ml |
| 7 | Recombinant Human Amphiregulin | Peprotech | 100-55B | 100 ng/ml |
| 8 | Recombinant Human Prolactin | Peprotech | 100-07 | 10 ng/ml |
| 9 | Human Insulin | Sigma | I2643- 250MG | 10 µg/ml |
| **Reagent B** |  |  |  |  |
|  | Hydrocortisone | Sigma | 1316004-200Mg | 0.5 µg/ml |

### PDXOs maintain estrogen receptor expression, are estrogen-responsive, and are sensitive to endocrine therapy

Consistent with ER expression detected in the PDX tumors *in vivo*, analysis of cell lysates from early (passage 2) and late passage (passage 20) organoid cultures expressed detectable ERalpha (Fig 2a), which did not change during long-term culture. Additionally, treatment of PDXOs with physiological levels of estrogen resulted in increased proliferation compared to vehicle control (Fig 2b). In contrast, culturing the organoids in phenol red-free media to exclude the weak estrogenic activity of phenol red induced a dramatic decrease in cell proliferation in PDX5 tumor-derived cultures that are known to express mutant ER (Guillen et al., 2022) (Fig 2c). These results are consistent with previous studies that showed these PDXs are responsive to estrogen stimulation (Gou et al., 2021; Guillen et al., 2022)Interestingly, organoid lines from PDX011 tumors expressing wild-type ER showed only a weaker but significant decrease in proliferation. Finally, to examine whether PDXOs can be used to study response to endocrine therapies, we treated PDXOs with the selective estrogen receptor degrader (SERD) fulvestrant, which is a widely used treatment for ER+ breast cancer. As expected, fulvestrant treatment resulted in a significant decrease in proliferation compared to vehicle control, suggesting that PDXOs, like ER+ breast tumors, require ER function to proliferate (Fig 2d). Thus, we demonstrate that ER+ PDXOs maintain ER expression, estrogen dependence, and responsiveness and retain endocrine sensitivity during long-term culture.

**Figure 2.**
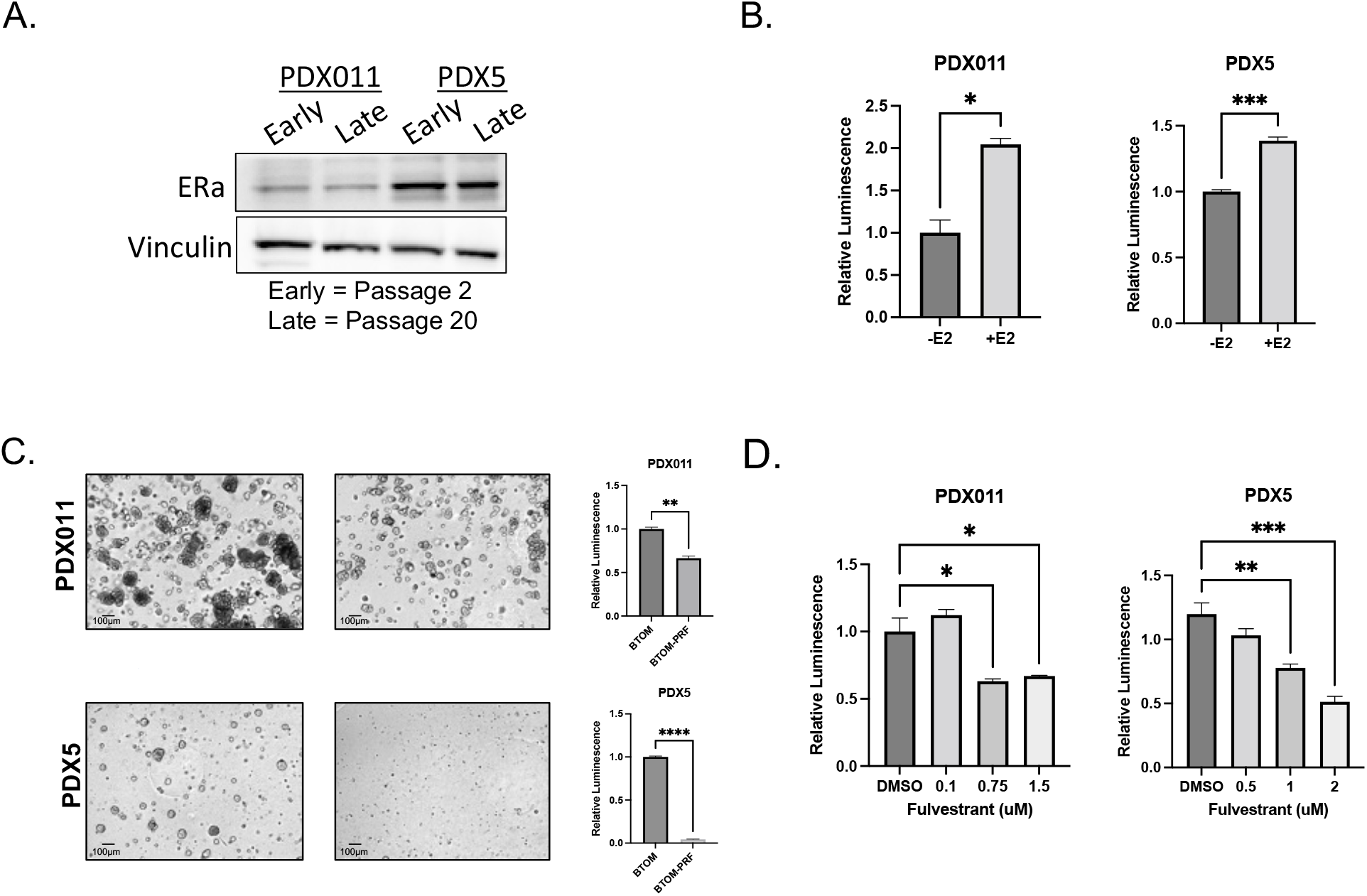
PDXOs grown in BTOM maintain estrogen-dependent phenotypes of ER+ breast cancer (A) Western blot showing Estrogen Receptor alpha expression in PDX011 and PDX5 organoids. VINCULIN was used as a loading control. (B) Luminescence levels in PDX011 and PDX5 organoids after 10 days of treatment with β-estradiol or vehicle control (EtOH). N=3. C) Representative brightfield images of Day 10 PDX011 and PDX5 grown in either BTOM or phenol red-free BTOM (left). Luminescence values for PDX011 and PDX5 organoids 10 days after being grown in either BTOM or phenol red-free BTOM (N=3). D) Luminescence values for PDX011 and PDX5 after being treated for 5 days with increasing doses of fulvestrant or vehicle control (N=3).

## Discussion

We report the development of an organoid culture media for the generation and maintenance of ER+ breast tumor organoids. We took into account growth factors and cytokines with established roles in ER-mediated signaling and growth, as well as maintenance of ER expression, leading to the inclusion of components such as prolactin, amphiregulin, IL6, and bovine pituitary extract. We excluded regulators of stemness, including WNT, R-Spondin-1, and Noggin, to avoid the selection of less differentiated epithelial lineages at the expense of differentiated lineages.

We demonstrate that ER+ organoids grown in BTOM-ER maintain the expression of estrogen receptor over 20 passages and maintain estrogen responsiveness. In addition, we demonstrate that the organoid lines remain sensitive to fulvestrant identifying BTOM-ER-generated primary tumor-derived organoid models can supplement the widely used ER+ cell lines, including MCF-7 and T47D, as a platform for investigating response and resistance to endocrine treatments. Given the consideration to growth factors that impact ER biology, it is likely BTOM-ER will be efficient in supporting the establishment and maintenance of ER+ luminal epithelial organoids from normal breast or mouse mammary glands; however, further studies will be required to investigate this possibility. Similar to many other tumor organoid platforms, the organoid culture conditions outlined here do not contain fibroblasts or immune cells, which have been shown to play a significant role in ER-mediated transcription and endocrine therapy response (Brechbuhl et al., 2017; Hiscox et al., 2006; Lang et al., 2013; Place et al., 2011). However, the culture conditions reported here are suitable for co-cultures, as we demonstrated recently for the culture of T cells with mouse or rat mammary tumor organoid models to investigate T-cell mediated killing of tumor epithelia (Gil Del Alcazar et al., 2022; Meng et al., 2021)to study the interaction between tumor-epithelial and stroma (Hogstrom et al., 2023). Thus, it is likely that the culture conditions reported here can serve as a new platform for understanding ER biology in primary breast tumor-derived epithelia that can lead to the development of new approaches to overcome drug resistance in ER+ breast cancer.

## Methods

### ER+ Breast Tumor Organoid Media (BTOM-ER)

For BTOM-ER growth media, 2.145 ml of Reagent A, 50 μl of Reagent B, and 1.0% penicillin-streptomycin were added to 100 ml of DMEM/F-12. Please see Table 1 for components of Reagent A and Reagent B and Supplemental Material 1 for further details on components and media recipes.

### Establishment of PDX organoid cultures

PDX tumor tissue was placed in a 10 cm dish, and 1-5 ml of resuspension media (DMEM-F12, % Penicillin-streptomycin, and 1.0 % BSA) was added to preserve the viability of the cells during processing. Tissue was then minced using sterile surgical scalpels into fragments ranging from 0.5-1.0 mm. Minced tissue was then transferred to a 15ml conical tube and pelleted by centrifugation at 1500 rpm for 5min at 4 degrees centigrade. After the supernatant was carefully removed, 2-10 ml of digestion media (DMEM-F12, 0.1mg/ml Collagenase/Dispase, and 1.0% Penicillin-streptomycin) was added, and the tube was placed in a 37 degrees centigrade shaker for 30-90 min, with checking every 15 min to check progress and ensure proper digestion of the tissue. The tissue is then pelleted again and resuspended in Accutase for 15-30 min to further digest tissue into organoid fragments. The tissue is then put through a 250 mm strainer to remove debris or large tissue fragments while allowing organoids to pass through. Organoids are pelleted and gently resuspended in BTOM-ER containing 5% Matrigel and ROCK-inhibitor (10 μM Y267632, Tocris), and then plated on top of a solidified Matrigel-coated well. Media was changed every 3-4 days.

### Passaging of PDXOs

After the removal of BTOM-ER, digestion media is added to each well, and the plate is incubated at 37 degrees centigrade for 1.5 hours. When the Matrigel has become a slurry and organoids are dispersed into small clumps or clusters, cold resuspension media is added at a 1:1 volume ratio to each well, and the suspension is transferred to a 15ml conical tube. After centrifugation at 1500 rpm for 5 min, the supernatant is removed, and the pellet is resuspended in Accutase and incubated for 20-30 min at 37 degrees centigrade. After incubation, resuspension media is added to the suspension. After centrifugation and removal of the supernatant, the pellet is resuspended in BTOM-ER and plated as done during the establishment of cultures.

### Immunohistochemistry (IHC) and Imaging of PDXOs

Immunohistochemistry was performed as previously described (Huang et al., 2020). Briefly, organoid tissue sections were generated by plating 20,000 cells/well of an 8-chamber slide. After the formation of mature organoid structures (typically 7-10 days post-plating), organoids were fixed in 4% PFA for 2 hours. After treatment with hematoxylin solution for 15 minutes, organoids were washed twice with water. The organoids were then scraped onto a solidified layer of histogel, with additional histogel added on top to create a sandwich within a cryomold. Once solidified, the histogel sandwich was transferred to a tissue cassette and fixed in 10% formalin for 16-24 h, followed by a brief wash in 70% EtOH. The cassettes were stored in 70% EtOH until submitted for sectioning. To obtain brightfield images of organoids in culture, cells were plated at a density of 25,000 cells/well, and images were taken at 20x, 10x, and 4x magnification using Spot Imaging software.

### Proliferation and Doubling Time of PDXOs

Organoids were digested as above and treated with TrypLE instead of Accutase to generate single cells. Each assay well in a 96-well plate was pre-coated with 30 ml of Matrigel and was allowed to solidify in a 37 degrees centigrade incubator for at least 10 min before plating cell suspensions on top. Cells were resuspended in BTOM-ER at a density of 50,000 cells/ml, and 100 ml was added to each well in triplicate. On the indicated days, corresponding wells were assessed for viability using 3D Cell Titer Glo (Promega). Viability measurements were normalized to Day 0, and doubling time was calculated using GraphPad Prism software.

### Western Blot Analysis

Organoids were grown in six-well plates at a density of 250,000 cells/ml. Once organoids reached confluency, each well was washed with ice-cold PBS. Matrigel was broken up by pipetting, and the slurry was transferred into a 15 ml conical tube. An additional 1.0 ml of PBS was added to the well to collect any remaining organoids. Organoids were pelleted by centrifugation at 3000 rpm for 5 min, and the supernatant was removed. The pellet was resuspended with 1.0 ml of Cell Recovery Solution (Corning) and incubated for one hour on ice. The released organoids were pelleted by centrifugation, and the supernatant was removed. After washing the pellet once with ice-cold PBS, the cells were resuspended with RIPA buffer, and western blotting analysis was performed. Signals were detected using Amersham Imager System via chemiluminescence.

### Estrogen Responsiveness and Dependence of PDXOs

For both estrogen responsiveness and dependence, organoids were digested and plated as above 40-50,000 cells/ml, depending on the organoid line. Three days after plating, media was refreshed, and wells were treated with either 1.0 nM of β−estradiol or vehicle control (EtOH) using the Tecan D300e drug dispenser. For dependence, media was replaced with fresh BTOM-ER or phenol red-free BTOM-ER. Media was refreshed every 2-3 days. After 10 days, organoids were assessed for viability using 3D Cell Titer Glo (Promega). Viability measurements were analyzed using GraphPad Prism software.

### Endocrine Therapy Treatment of PDXOs

Organoids were prepared as above, and on day three, media was refreshed, and wells were treated with either vehicle control or indicated concentrations of fulvestrant (Selleckchem) using the Tecan D300e drug dispenser. Media was refreshed, and plates were retreated every 2-3 days. After 5 days of treatment, organoids were assessed for viability using 3D Cell Titer Glo (Promega).

### Statistics and reproducibility

Error bars were generated by Standard Error Mean (SEM) calculations. For experiments with two conditions, an unpaired one-tailed Student’s T-test was performed. For experiments with three or more conditions, one-way ANOVA followed by a Bonferroni comparison was used. Doubling time was calculated by applying a non-linear regression for exponential growth for each condition.

## Acknowledgments

We thank the PDXNet Consortium, especially Dr. Alana Welm and Dr. Michael T. Lewis, for providing the PDX models used for this study. The authors would also like to thank the Beth Israel Deaconess Medical Center Histology Core for sectioning and staining of the PDXO samples. Lastly, the authors would like to thank Ridhdhi Desai, Weilin Li, Ling Huang, and Almudena de Gregorio for their technical assistance, constructive feedback, and helpful discussions. This work was supported by the National Institutes of Health grant 5K00CA223023 (MUJO), the Breast Cancer Research Foundation (SKM), and the NCI Intramural program (SKM).

