## Supplemental materials for "Establishing conditions for the generation and maintenance of estrogen receptor-positive organoid models of breast cancer"

**Suppl. Fig. 1**

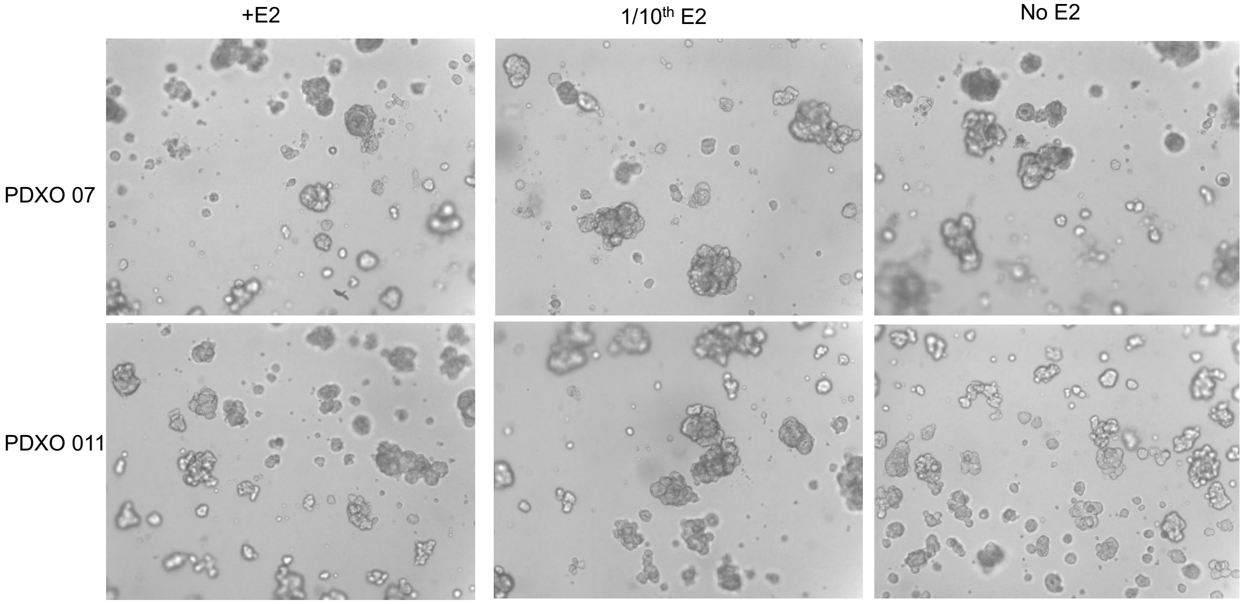


**Suppl. Fig. 1: Effect of β-estradiol supplementation to growth media.**

Two organoid lines were grown in the presence of E2 (1.0 nM β-estradiol) or 1/10th the concentration or no E2, and morphology was monitored by analysis of phase images over 12 days. A representative image from day 7 of the culture is shown.

### Supplemental Material 1: Detailed methods for preparation of ER+ breast tumor organoid media

The initial organoid culture that develops after the plating of the processed tissue should be referred to as Passage 0 (P0). All subsequent passages should be serially numbered, and records maintained for each passage. We recommend freezing down early passage cultures before using them for experiments. We typically do not use cultures older than P20 for experiments for concerns related to culture-induced genetic drift.

- *We prepare master cocktails pre-mix and store them in small aliquots for extended periods of time (6 months).*
- *The master cocktail pre-mix is diluted in cell culture media with antibiotics and stored at 4 degrees for short-term use (~1 month).*
- *Just before every use, we prepare 'growth media' that contains 5% Matrigel.*

**Materials Needed**

| **Materials** | **Vendor** | **Catalog #** |
|---|---|---|
| Collagenase/Dispase(100 mg/ml) | Sigma | 11097113001 |
| Accutase | Sigma | A6964 |
| DMEM+F-12 | Thermo | 11330-032 |
| Rock Inhibitor (Y267632) | Tocris | 1254 |
| Matrigel (Growth factor reduced) (Concentration ~8.5mg/ml) | BD | 354230 |
| Penicillin-Streptomycin 10,000 U/mL | Gibco | 15140-122 |
| Bovine Serum Albumin Heat shock fraction | Sigma | A7906 |
| Growth Media | See recipe below | |

| | | |
|---|---|---|
| BTOM | See recipe below | |
| Tissue strainer (250µm) | Thermo | 87791 |
| Chamber slides | BD or any vendor | |
| Freezing media (Cryostar cs10) | Stem Cell Tech | 7930 |

**BTOM-ER Pre-mix cocktail recipe**

| | Supplements/Growth factors | Vendor | Catalog # | Final concentration |
|---|---|---|---|---|
| **Reagent A – (**See note below) | | | | |
| 1 | BPE | Hammond cell tech | 1078-NZ | 0.8 ml for 100ml |
| 2 | B27 | Thermo | 17504001 | 1.0 ml for 100ml |
| 3 | Recombinant Human FGF- Basic (FGF2) | Peprotech | AF-100- 18B | 10ng/ml |
| 4 | Recombinant Human FGF10 | Peprotech | 100-26 | 10ng/ml |
| 5 | Recombinant Human EGF | Peprotech | AF-100-15 | 2ng/ml |
| 6 | Recombinant Human IL6 | Peprotech | 200-06 | 100ng/ml |
| 7 | Recombinant Human Amphiregulin | Peprotech | 100-55B | 100ng/ml |
| 8 | Recombinant Human Prolactin | Peprotech | 100-07 | 10ng/ml |
| 9 | Human Insulin | Sigma | I2643- 250MG | 10ug/ml |
| **Reagent B – (**See note below) | | | | |
| | Hydrocortisone | Sigma | 1316004-200Mg | 0.5ug/ml |

### Notes:

Aliquoting B27.

1. Place the frozen B27 stock in a 4-degree fridge overnight.

2. On the second day, gently invert the fully thawed B27 a few times and aliquot into 2.0 ml aliquots.
3. Store the tubes in a -80 degree freezer.

<u>Aliquoting BPE</u>

1. Place the frozen BPE in a 4-degree fridge overnight to thaw.
2. Incubate the fully thawed BPE in a 37-degree water bath for one hour. This helps lipids in BPE to dissolve.
3. Transfer the 100 mL BPE into two 50 mL conical tubes and centrifuge at 2500 rpm for 5 minutes.
4. Use a 10 mL pipette to transfer the supernatant into two new 50 mL conical tubes. Discard the precipitates.
5. Aliquot BPE into 2.0 ml microcentrifuge tubes and store in a -80 freezer.

<u>Growth Factors:</u>
Prepare growth factor stock solutions as 1000x stocks following per manufacturer's recommendation and store them in 100 ml aliquots.

<u>Hydrocortisone:</u>

Prepare a 1000x stock and store it in 100 µl aliquots.

### <u>Media Recipes:</u>

**Digestion Media:**
DMEM/F12
1:100 dilution of 100mg/ml stock Collagenase/ Dispase
1.0 % Penicillin-streptomycin
<u>Preparation Tip:</u> Prepare only the needed amount, fresh just before use, and discard the unused portion of the media.

**Resuspension Media**:
DMEM/F12
1.0 % BSA
1.0 % Penicillin-Streptomycin
<u>Preparation Tip:</u> Add 1% BSA by weight to DMEM+1.0% Penicillin- streptomycin-containing media and stir to dissolve the BSA. Once dissolved, filter sterilize using a 0.2µ filter and store at 4°C.

**BTOM-ER Growth Media:**
DMEM/F-12: 100 ml
2.145 ml of Reagent A

50 µl of Reagent B

1.0% Penicillin-Streptomycin

Preparation Tip:

*Do NOT mix Reagent A and Reagents B by themselves as the alcohol in Reagent B will denature the growth factors*

After adding all ingredients, filter the media through a 0.2µ filter

Store prepared media at $4^0$C for NO more than one month.

#### Culture Media (make fresh for immediate use):

BTOM-ER growth media.

5.0 % GFR-Matrigel

10 µM Y267632, Rock inhibitor (10mM, a 1000x stock). Rock inhibitor is prepared in sterile Phosphate Buffered Saline.

Preparation Tip: Prepare fresh. You may premix BTOM-ER growth media and ROCK inhibitor and keep them on ice. Add Matrigel just before use. Discard unused portions of the media.

#### Freezing Media:

Cryostar freezing media

10 µM Y267632, Rock inhibitor
